# *Chlamydia trachomatis* induces the transcriptional activity of host YAP in a Hippo-independent fashion

**DOI:** 10.1101/2022.03.16.484654

**Authors:** Liam T. Caven, Amanda J. Brinkworth, Rey T. Carabeo

## Abstract

The obligate intracellular pathogen *Chlamydia trachomatis* is the causative agent of the most common bacterial sexually transmitted disease worldwide. While the host response to infection by this pathogen has been well characterized, it remains unclear to what extent host gene expression during infection is the product of *Chlamydia*-directed modulation of host transcription factors. In this report, we show the transcriptome of *Chlamydia*-infected epithelial cells exhibits gene expression consistent with activity of YAP, a transcriptional coactivator implicated in cell proliferation, organ morphogenesis, wound healing, and fibrosis. After confirming induction of YAP target genes during infection, we observed increased YAP nuclear translocation in *Chlamydia*-infected epithelial cells. We show that increased YAP activation is a *Chlamydia*-directed process occurring during midcycle infection; critically, this phenotype is sensitive to inhibition of protein synthesis by the pathogen. Infection-mediated YAP activation bypasses YAP inhibition by the Hippo kinase cascade, instead involving YAP tyrosine phospho-activation and the activity of host Src family kinases. Taken together, our results define a mechanism for *Chlamydia*-directed modulation of host gene expression independent of the host inflammatory response to infection, as well as introduce novel therapeutic targets for treatment of this pervasive disease.

## 1 Introduction

*Chlamydia trachomatis* is the most common bacterial cause of both sexually-transmitted disease and infectious blindness worldwide. An obligate intracellular pathogen, *C. trachomatis* binds to and induces its own internalization by epithelial cells, establishing a replicative niche within a pathogen-remodeled vacuole termed the chlamydial inclusion. Chronic or repeated *C. trachomatis* infection is associated with severe inflammatory and fibrotic sequelae. *C. trachomatis* serovars A-C, which preferentially infect the conjunctiva, are associated with excess collagen production and contraction of infected tissues, leading to the inward turning of the upper eyelid and subsequent corneal abrasion by the eyelashes (Taylor et al., 2014; Lansingh, 2016). *Chlamydia*-mediated trachoma is endemic to Africa, Latin America, Asia, and the Middle East (Hu et al., 2010), and is responsible for the blindness or visual impairment of up to 1.9 million people according to current WHO estimates. Infection by *C. trachomatis* serovars D-K, which preferentially colonize the reproductive tract, are associated with pelvic inflammatory disease as well as progressive scarring of the fallopian tubes (Ness et al., 2008; Haggerty et al., 2010a). Blockage of the upper female genital tract by scar tissue can consequently lead to ectopic pregnancy or tubal factor infertility (Chow et al., 1990). In the United Kingdom, *C. trachomatis* is estimated to account for 29% of all cases of tubal factor infertility, which in turn represents 25% of all cases of female infertility (Price et al., 2016). Collectively, the urgent threat to public health presented *Chlamydia*-associated sequelae has made understanding the underlying mechanisms of the host-*Chlamydia* interaction a subject of ongoing investigation.

Initial study of the host response to infection has made a persuasive argument for infection-associated pathology being the product of cytokine signaling cascade originating in infected epithelial cells, termed the cellular paradigm of pathogenesis (Stephens, 2003). Clinical infections have revealed substantial infiltration of infected tissues by immune cells *in vivo*, suggesting that infected epithelial cells recruit components of the innate and adaptive immune systems to the site of infection (Kiviat et al., 1990; Stephens, 2003). Indeed, upregulation of an expansive portfolio of inflammatory cytokines has been consistently observed in chlamydial infections *in vitro*, including GROα, GM-CSF, IL-1α, IL-6, IL-11, and IL-8 (Eckmann et al., 1993; Rasmussen et al., 1997; Cheng et al., 2008). However, the pro-inflammatory host response fails to account for the fibrotic outcomes of asymptomatic infection; it is estimated that up to 75% of *C. trachomatis* infections go unreported due to absent or subclinical symptoms, and that up to 18% of such cases are believed to cause infertility (Haggerty et al., 2010b).

One explanation for this discrepancy in the data is that *C. trachomatis* induces pathology via direct manipulation of host gene expression; indeed, mounting evidence suggests that regulation of host transcription is targeted by the pathogen in a variety of ways. Past work has shown that *C. trachomatis* antagonizes pro-inflammatory signaling mediated by the transcription factor NF-κB by stabilizing its inhibitory subunit IκBα, through the activity of the chlamydial deubiquitnase ChlaDub1 (Le Negrate et al., 2008). The chlamydial proteinase CPAF has also been shown to inhibit the NF-κB complex, in that ectopic expression of this chlamydial effector leads to degradation of the complex subunit p65/RelA (Christian et al., 2010). However, the specific role of CPAF in inhibiting pro-inflammatory gene expression remains unclear, given that infection with CPAF-deficient mutant strains does not exhibit increased NF-κB activation (Snavely et al., 2014), likely due to compensatory activity of ChlaDub1 and other potential mechanisms of chlamydial NF-κB antagonism. In similar fashion, infection with the related species *C. pneumoniae* has been shown to induce expression and phosphorylation of the AP-1 transcription factor c-Jun (Krämer et al., 2015). Intriguingly, this effect did not occur during infection with heat-killed *C. pneumoniae*, indicating that c-Jun activation is a pathogen-directed phenotype with implications for virulence, given that c-Jun knockdown negatively impacted bacterial load as well. Collectively, these data illustrate the potential for *Chlamydiae* to induce changes in host gene expression to facilitate pathogenesis, demonstrating a need to better characterize how the pathogen acts on host transcription factors.

To that end, we have examined the host transcriptome of *C. trachomatis-infected* immortalized endocervical epithelial cells, to perform unbiased *in silico* discovery of pathogen-modulated host transcription factors. In this report, we show that *C. trachomatis* infection induces gene expression consistent with the function of YAP (<u>Y</u>es-<u>a</u>ssociated <u>p</u>rotein), a transcriptional coactivator. After empirically confirming infection- and YAP-dependent induction of target genes identified by this analysis, we subsequently demonstrate that infection promotes YAP activity by enhancing its nuclear translocation. Increased YAP nuclear translocation was first detectable at 18 hours post-infection and exhibited sensitivity to inhibition of bacterial protein synthesis, indicating that YAP induction is a pathogen-directed process. Intriguingly, chlamydial activation of YAP occurs irrespective of the Hippo kinase cascade (a well characterized inhibitor of YAP activity), instead relying upon YAP phosphorylation at Y357, a post-translational modification shown to enhance YAP activation (Smoot et al., 2018; Sugihara et al., 2018). We show that phosphorylation at this residue is essential for infection-mediated YAP nuclear translocation. Additionally, we find that enhanced YAP nuclear translocation and Y357 phosphorylation in infected cells is sensitive to inhibition of Src-family kinases, consistent with prior reports implicating this kinase family in Hippo-independent YAP activation. With these data, we present an initial framework by which *Chlamydia trachomatis* manipulates the activity of a transcription factor to drive changes in host gene expression.

## 2 Materials and Methods

### 2.1 Eukaryotic Cell Culture

Human endocervical epithelial HPV-16 E6/E7 transformed End1s (End1 E6/E7, ATCC CRL-2615) were cultured at 37° C with 5% atmospheric CO_2_ in Keratinocyte Serum-Free Medium (KSFM, Thermo Fisher Scientific) supplemented with human recombinant epidermal growth factor, bovine pituitary extract, 5 micrograms/mL gentamicin, and 0.4 mM CaCl_2_ (unless otherwise indicated). For all experiments, End1s were cultured between passages 3 and 15. Primary human cervical epithelial cells (HCECs, ATCC PCS-0480-011, Lot 80306190) were cultured at 37° C with 5% atmospheric CO_2_ in Cervical Epithelial Cell Basal Medium (CECBM, ATCC PCS-480-032) supplemented with all contents of a Cervical Epithelial Growth Kit (ATCC PCS-080-042). Raft cultures were prepared as previously described by our laboratory (Nogueira et al., 2017), exchanging HaCaT cells for End1s where indicated, and cultured over 19 days before being fixed for 30 minutes in 4% paraformaldehyde, OCT-embedded, cryo-sectioned, and stained with hematoxylin and eosin.

### 2.2 Cloning and DNA/siRNA Transfection

Both FLAG-YAP1-WT and FLAG-YAP1-Y357F were generated via the simplified FastCloning method of Gibson assembly (Li et al., 2011); briefly, PCR-amplified recombinant DNA was treated with DpnI (NEB Biolabs) for 3 h, then transformed into Stellar chemically competent *E. coli* (Clontech 636763) via heat shock, followed by Kanamycin selection, screening via colony PCR (where applicable), and sequence confirmation. FLAG-YAP1-WT was generated via amplification of pEGFP-C3-HYAP1, a gift from Marius Sudol (Addgene plasmid # 17843; http://n2t.net/addgene:17843; RRID:Addgene_17843), using the following primers: FWD: 5’-GACTACAAAGACGATGACGACAAGGATCCCGGGCAGCAGCCG-3’; REV: 5’-CTTGTCGTCATCGTCTTTGTAGTCCATGGTGGCGACCGGTAGCG-3 ‘. FLAG-YAP 1-Y357F was generated from FLAG-YAP1-WT via site-directed mutagenesis of the Y357 codon (TAC to TTC), using the following primers: FWD: 5’-GCAGCTTCAGTGTCCCTCGA-3’, REV: 5’-ACACTGAAGCTGCTCATGCTTAGTC-3 ‘.

End1s were transfected with plasmid DNA via electroporation: after harvest via trypsinization, 2×10^6^ Endls per construct were resuspended in 400 μL Opti-MEM at room temperature (Thermo Fisher Scientific 31985062) containing 20 ug of plasmid DNA and 3 μL sheared salmon sperm DNA (Thermo Fisher Scientific AM9680). Suspensions were then transferred to individual GenePulser 0.4 cm cuvettes (Bio-Rad 1652088), then electroporated using a GenePulser XCell (Bio-Rad) using the following parameters: exponential decay program template, 225 V, 850 μF capacitance, infinite resistance. 100 μL aliquots of the resulting electroporants were transferred to individual wells of a 24-well plate containing cover slips and pre-warmed KSFM, then incubated until confluent monolayers had formed (24-48h) for subsequent infection as previously described.

For siRNA-mediated knockdown experiments, End1s were transfected with either an ON-TARGETplus non-targeting siRNA pool (Horizon Discovery D-001810-10-05) or an ON-TARGETplus YAP1-targeting siRNA SMARTpool (Horizon Discovery L-012200-00-0005) using Lipofectamine 3000 (Thermo Fisher Scientific L3000008), per manufacturer’s instructions, at an empirically determined optimal concentration of 10 nM. At 16 h post-seeding of End1s at 125% of confluence on 6-well plates as described above, siRNA was combined in Opti-MEM (Thermo Fisher Scientific 31985062) with the Lipofectamine 3000 reagent, incubated for 5 m at room temperature to allow for liposome formation, then added to wells dropwise with mixing. Transfected End1s were then incubated for 24 h prior to infection with *Chlamydia* (see below).

### 2.3 Chlamydial Infections

*Chlamydia trachomatis* serovar L2 (434/Bu) was originally obtained from Dr. Ted Hackstadt (Rocky Mountain National Laboratory, NIAID). Chlamydial EBs were isolated from infected, cycloheximide-treated McCoy cells at 36-40 hours post-infection (hpi) and purified by density gradient centrifugation as previously described (Caldwell et al., 1981). For infection of 6- and 24-well tissue culture plates (Greiner Bio-One 657160 and 662160), End1s seeded at 125% of confluence (16 h post-seeding) or HCECs grown to confluence (6-7 days) were washed with pre-warmed Hanks Buffered Saline Solution (HBSS) prior to inoculation with *Chlamydia*-containing KSFM or CECBM at a multiplicity of infection (MOI) of 5, unless otherwise indicated. Tissue culture plates were centrifuged at 4° C and 500 rcf (Eppendorf 5810 R tabletop centrifuge, A-4-81 rotor) for 15 minutes to synchronize infection. Inoculum was then aspirated, and cells were washed with chilled HBSS prior to the addition of pre-warmed KSFM or CECBM.

### 2.4 Bulk RNA-Sequencing and Analysis

End1s seeded on fibronectin-coated 6-well plates (Corning 354402) and infected at an MOI of 2 as described above were harvested for RNA using TRIzol (Thermo Fisher Scientific 15596026) and the DNA-free DNA removal kit (Thermo Fisher Scientific AM1906), according to manufacturers’ protocols. Extracted RNA was subsequently enriched for polyadenylated mRNA transcripts using the NucleoTrap mRNA enrichment kit (Macherey-Nagel) according to the manufacturer’s protocol. For the comparison of mock- and *Ct* serovar L2-infected transcriptomes, RNA samples were subsequently assayed for fragmentation using an Agilent 5200 Fragment Analyzer; intact samples were enriched for polyadenylated transcripts via the NucleoTrap kit a second time due to the high incidence of ribosomal RNA in the fragment analysis results. cDNA library preparation was performed using the Ion Xpress Plus Fragment Library kit (Thermo Fisher Scientific 4471269), and sequencing of cDNA libraries was performed using the Ion Torrent system (Life Technologies). For the bulk RNA-sequencing of primary cells, HCECs infected at an MOI of 2 were harvested for RNA samples via TRIzol and the DNA-free DNA removal kit as described above, and were subsequently assayed for fragmentation using an Agilent Bioanalyzer 2100. cDNA library preparation of intact samples was performed using the NuGEN Universal mRNA-Seq Library Preparation kit, and sequencing of cDNA libraries was performed using the NextSeq 550 system (Illumina).

Read alignment and downstream analysis of both experiments was performed using CLC Genomics Workbench (Qiagen); each treatment group was comprised of libraries from three biological replicates, each with a minimum of 30 million reads (unstranded single read, mean length 150 bp), genes with an FDRP ≤ 0.05 were considered differentially expressed. ChEA crossreferencing was performed in Python, identifying the number of genes induced (FC ≥ 1.5, FDRP ≤ 0.05) and repressed (FC ≤ 1.5, FDRP ≤ 0.05) by infection for each transcription factor’s gene targets (Lachmann et al., 2010). Pearson’s correlation coefficients between End1 and HCEC expression of YAP-responsive genes were subsequently calculated in R, using log_2_-transformed fold changes of all genes differentially expressed in at least one data set.

### 2.5 Reverse Transcription Quantitative Real-Time PCR

End1s seeded on fibronectin-coated 6-well plates (Corning 354402) and infected or cocultured as described above were harvested for RNA using TRIzol (Thermo Fisher Scientific 15596026) and the DNA-free DNA removal kit (Thermo Fisher Scientific AM1906), according to manufacturers’ protocols. cDNA libraries were subsequently prepared using SuperScript IV Reverse Transcriptase (Thermo Fisher Scientific 11766050) according to the manufacturer’s protocol. Quantitative real-time PCR was performed on a QuantStudio 3 (Thermo Fisher Scientific) using TaqMan assay kits (Thermo Fisher Scientific) of the following genes: CTGF (Hs01026927_g1), INHBA (Hs01081598_m1), BMP2 (Hs00154192_m1), MMP9 (Hs00957562_m1), and the housekeeping gene HPRT (Hs02800695_m1). Statistical analysis was performed in R, using pairwise Student’s t-tests and Bonferroni’s correction for multiple comparisons; p-values less than 0.05 were considered statistically significant.

### 2.6 Immunofluorescence Microscopy

End1s or HCECs seeded on glass cover slips (VWR) coated with fibronectin (Corning 354008) or fibronectin-deposited micropatterned chips (CYTOO) and infected as described above were fixed in 4% paraformaldehyde in phosphate-buffered saline (PBS) for 10 minutes at 37° C, washed in PBS, then blocked in 5% bovine serum albumin (BSA) in PBS for 1 hour at room temperature. Fixed and blocked cover slips were subsequently incubated overnight at 4° C with primary antibodies in 1% BSA-PBS: rabbit anti-YAP (CST 4912, 1:100 dilution), rabbit anti-E-Cadherin (CST 3195, 1:200 dilution). Cover slips were again washed in PBS, then incubated for 1 hour at room temperature with the following fluorophore-conjugated antibodies/dyes in 1% BSA-PBS: goat anti-rabbit Alexa-488 conjugate (Thermo Fisher Scientific A-11034, 1:1000 dilution), phalloidin Alexa-594 conjugate (Thermo Fisher Scientific A-12381, 1:120 dilution), DAPI (Sigma-Aldrich 10236276001, 1:1000 dilution). Afterward, cover slips were washed in PBS and ultrapure water, then mounted on microscope slides using Shandon Immu-Mount (ThermoFisher Scientific 9990402).

A minimum of 5 fields per cover slip were imaged using a SP-8 Lightning Confocal Microscope (Leica) or CSU-W1 Spinning-Disk Confocal Microscope (Nikon). To account for variations in nuclear size and avoid biased selection of cells for measurement, blinded image quantification was performed by assigning image filenames randomized number codes, selecting 10 nuclei at random per field using only the DAPI channel, manually masking the nuclear area, and recording the mean YAP fluorescence intensity per nucleus. To account for variation in total YAP between cells, staining efficiency between cover slips, and compression of the nuclear/cytosolic compartments by the chlamydial inclusion, 5 cytosolic regions not occluded by an inclusion body and adjacent to measured nuclei were selected per field, with the mean YAP fluorescence intensity of these regions averaged to produce a per-field measurement of mean cytosolic YAP fluorescence intensity; nuclear translocation of YAP was thereby expressed as a ratio of mean nuclear fluorescence to mean cytosolic fluorescence. Statistical analysis was performed in R, using a Kruskal-Wallis test to first verify a statistically significant (p-value < 0.05) difference between treatment groups. Subsequent pairwise comparisons were performed using a Wilcoxon rank sum test and Bonferroni’s correction for multiple comparisons, with p-values less than 0.05 being considered statistically significant.

### 2.7 SDS-PAGE and Western Blotting

To minimize activity of the chlamydial protease CPAF, End1s seeded on 6-well plates and infected as described above were subsequently lysed in 1% SDS buffer heated to 95° C, as previously described (Johnson et al., 2015). After treatment with Pierce Universal Nuclease (Thermo Fisher Scientific, 1:1000 dilution) for 5 minutes at room temperature, lysates were combined with 4X Laemmli Sample Buffer (Bio-Rad 1610747) for loading on a 7.5% acrylamide/bis-acrylamide gel for SDS-PAGE (1.5 hours, 120V). Gels were then transferred to PVDF membranes (Bio-Rad 1620177) using a semi-dry transfer method. After blocking in 5% BSA in PBST (PBS containing 0.1% Tween-20) for 1 hour at room temperature, membranes were incubated overnight at 4° C with primary antibodies in 5% BSA-PBST: rabbit anti-YAP (CST 4912, 1:1000 dilution), rabbit anti-S127-pYAP (CST 13008, 1:1000 dilution), rabbit anti-Y357-pYAP (Abcam ab62571, 1:1000 dilution). Membranes were subsequently washed in PBST, then incubated with a goat anti-rabbit HRP-conjugated secondary antibody (Dako P0448, 1:2000 dilution in 5% BSA-PBST) for 2 hours at room temperature. After additional washing in PBST, membranes were imaged using Immobilon HRP Substrate (Millipore Sigma WBKLS0500) or an Azure Biosystems c600. Images were analyzed using the ImageJ gel analysis tool to quantify the fluorescence density of phospho-analyte bands relative to the YAP total protein loading control. Statistical analysis was performed in R, using pairwise Student’s t-tests with Bonferroni’s correction for multiple comparisons; p-values less than 0.05 were considered statistically significant.

## 3 Results

### 3.1 *Chlamydia* infection induces the expression of a subset of YAP target genes

To identify host transcription factors modulated by *Chlamydia* infection, we opted to model infection of the upper female genital tract using End1/E6E7 immortalized, non-transformed endocervical epithelial cells (Fichorova et al., 1997). We first confirmed the non-tumorigenic growth properties of End1 cells via organotypic raft culture: first seeding End1 cells on collagen gels in a transwell insert, then inducing differentiation via exposure to the air-liquid interface (Nogueira et al., 2017). Transformed cervical epithelial cells form irregular layers when induced to differentiate via air-liquid interface (Aasen et al., 2003); by contrast, differentiating End1s grown via raft culture formed epithelial tissues of uniform thickness (Supplementary Figure S1), consistent with previous reports of organotypic culture with this cell line (Gali et al., 2010). Having thus validated our experimental model, we then infected End1s with the lymphogranuloma venereum *Chlamydia trachomatis* serovar L2, then harvested RNA for bulk RNA-sequencing at 24 hours post-infection (hpi). This timepoint occurs well after chlamydial differentiation into proliferative and metabolically active reticulate bodies (Lee et al., 2018), thus allowing for production of any chlamydial factors modulating host gene expression. Critically, infection induced the differential expression (FDRP ≤ 0.05) of 3611 genes relative to a mock-infected control (Figure 1A, Supplementary Data S1, GEO Accession: GSE180784).

**Figure 1:**
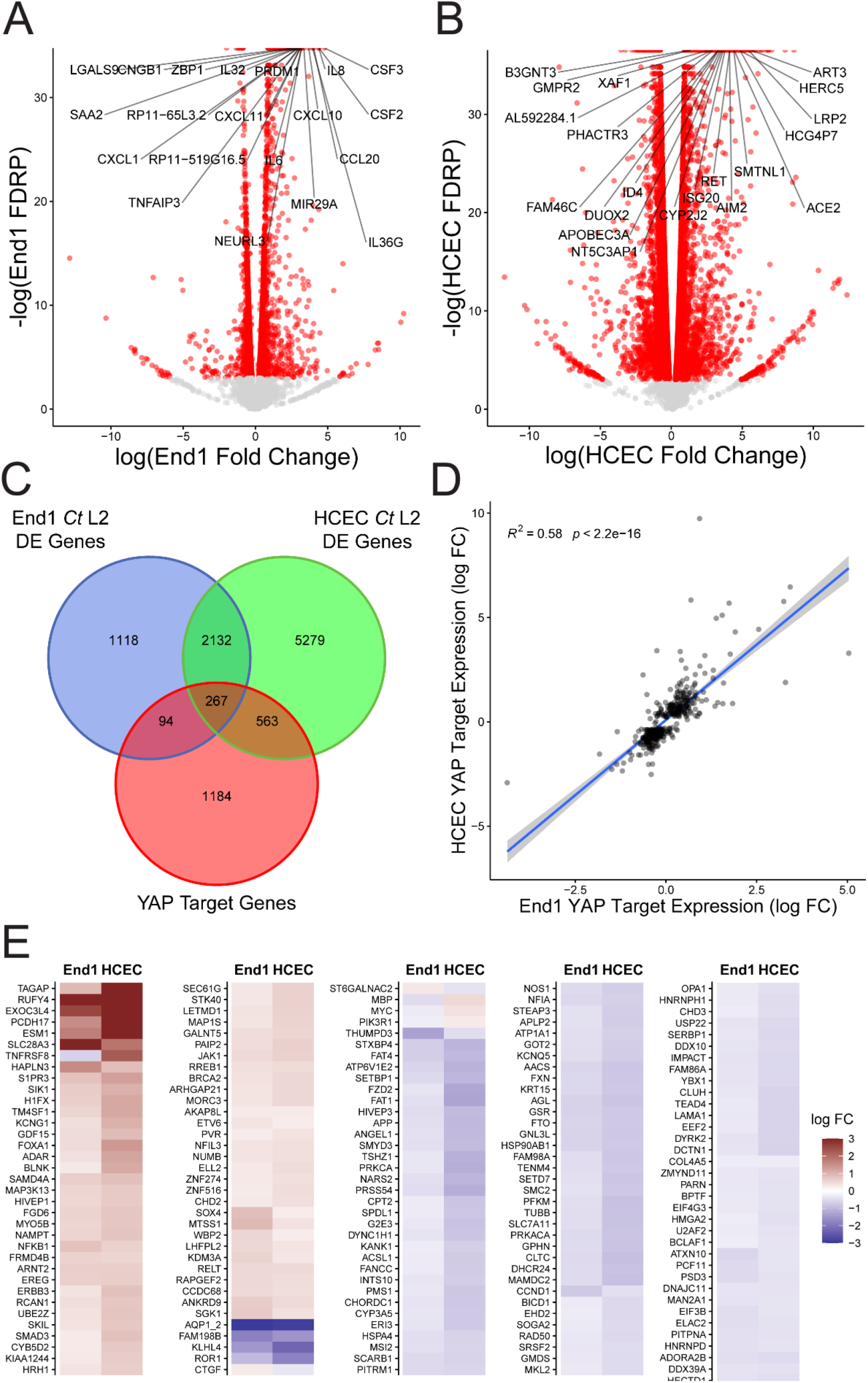
*Chlamydia* infection induces expression of a subset of YAP target genes. (A, B) Volcano plot of gene expression changes identified via bulk RNA-sequencing of End1/E6E7 immortalized epithelial cells (A) and primary human endocervical epithelial cells (B) during infection with *Chlamydia trachomatis* (*Ct*) serovar L2 compared to mock infection, at 24 hours post-infection (hpi). n = 3, with a minimum of 3×10 unstranded single reads per replicate with a mean length of 150 bp. All fold changes are relative to the mock-infected control; red dots: false discovery rate p-value (FDRP) ≤ 0.05, labels: top 20 genes whose expression differed most significantly (lowest FDRP) from to the mock infection. (C)Venn diagram of differentially expressed genes identified in (A) and (B) cross-referenced with the ChIP Enrichment Analysis (ChEA) database of YAP target genes. (D)Scatter plot of gene expression of YAP-responsive (ChEA), differentially expressed genes in either *Ct* serovar L2-infected End1/E6E7 cells (End1s, x-axis) or primary human cervical epithelial cells (HCECs, y-axis). All fold changes are relative to each cell type’s respective mock-infected control; blue line: linear regression model of correlation; grey shading: 95% confidence interval. R^2^ and p-values calculated using Pearson’s correlation. (E)Heatmap of YAP target gene expression in *Ct* serovar L2-infected End1s (left columns) and HCECs (right columns). All fold changes are relative to each cell type’s respective mock-infected control; only genes differentially expressed (FDRP ≤ 0.05) in both cell types are shown.

To then determine potential transcription factors driving the host response to infection, we then cross-referenced this differentially expressed gene set with the ChEA database of transcription factor gene targets, derived from gene set enrichment analysis of published ChIP-chip, ChIP-seq, ChIP-PET, and DamID data (Lachmann et al., 2010). Of the 199 transcription factors included in ChEA, 149 exhibited twice as many induced target genes (FC ɥ 1.5) as repressed target genes (FC ≤ 1.5), suggesting chlamydial induction of transcription factor activity (Supplementary Data S2). Critically, targets of transcription factors known to regulate inflammation (IRF1, STAT1, STAT3) and fibrosis (SMAD2/3, EGR1, TEAD4, YAP1) appeared to be highly induced in the host (Table 1), consistent with prior reports of infection-dependent induction of pro-inflammatory and pro-fibrotic gene expression (Carlson et al., 2005; Lad et al., 2005; Humphrys et al., 2013; Sun et al., 2017).

**Table 1:** Cross-referencing of RNA-sequencing results with ChEA transcription factor target database reveals induction of transcription factor targets.

| <u>Transcription Factor</u> | <u>Induced Targets</u> | <u>Repressed Targets</u> | <u>Mean Fold Change</u> |
| --- | --- | --- | --- |
| IRF1 | 55 | 0 | 2.6 |
| STAT1 | 34 | 9 | 1.44 |
| STAT3 | 255 | 80 | 2.46 |
| SMAD2 | 126 | 31 | 9.11 |
| SMAD3 | 162 | 57 | 4.89 |
| TEAD4 | 67 | 29 | 1.21 |
| EGR1 | 292 | 101 | 2.11 |
| YAP1 | 81 | 36 | 1.24 |
| MYCN | 61 | 56 | -62.65 |
| MYBL1 | 2 | 4 | -0.49 |
| CRX | 21 | 18 | -2.02 |

Intriguingly, the fibrosis-associated transcription factors identified by this analysis (SMAD2/3, EGR1, and TEAD4) are known binding partners of the transcriptional coactivator YAP1 (Zagurovskaya et al., 2009; Zanconato et al., 2015; Szeto et al., 2016). While YAP1 target gene induction was modest compared to other transcription factors identified by this method, the more robust target gene induction of transcription factors upregulated by YAP suggests the potential for chlamydial modulation of this transcriptional coactivator. To assess the physiological relevance of apparent YAP-associated gene expression, we then sequenced polyadenylated RNA from primary human endocervical epithelial cells (HCECs) infected with *Ct* serovar L2 (Supplementary Data S3, GEO Accession: GSE180784). Infected HCECs exhibited differential expression (FDRP ≤ 0.05) of 8241 genes (Figure 1B), of which 839 mapped to the ChEA database of YAP-responsive genes (Figure 1C). Expression of YAP-responsive genes by infected End1s and HCECs exhibited significant correlation (Figure 1D), suggesting that infection of these cell types activates a conserved, putatively YAP-mediated transcriptional program. Intriguingly, only a subset of YAP target genes is upregulated in either End1s or HCECs, suggesting that *Chlamydia* may alter the YAP regulon in host cells, or potentially modulate YAP transcriptional activity (Figure 1E). Taken together, these data imply that the transcriptome of *Chlamydia-infected* epithelial cells may be in part be driven by YAP activity.

### 3.2 *Chlamydia* infection promotes YAP nuclear translocation

To confirm our RNA-sequencing analysis suggesting infection may promote YAP-dependent gene expression, we next assayed expression of connective tissue growth factor (CTGF) – a consistent marker of YAP target gene induction (Futakuchi et al., 2018). We observed host CTGF expression was significantly enhanced by infection, as was expression of additional genes known to be YAP-responsive (Figure 2A), including inhibin beta A (INHBA), bone morphogenetic protein 2 (BMP2), and matrix metallopeptidase 9 (MMP9) (Mo et al., 2012; Huang et al., 2016; Pan et al., 2017). Critically, CTGF expression was sensitive to siRNA-mediated YAP knockdown (Figure 2B, Supplementary Figure S1), suggesting that infection modulates gene expression in a YAP-dependent fashion

**Figure 2:**
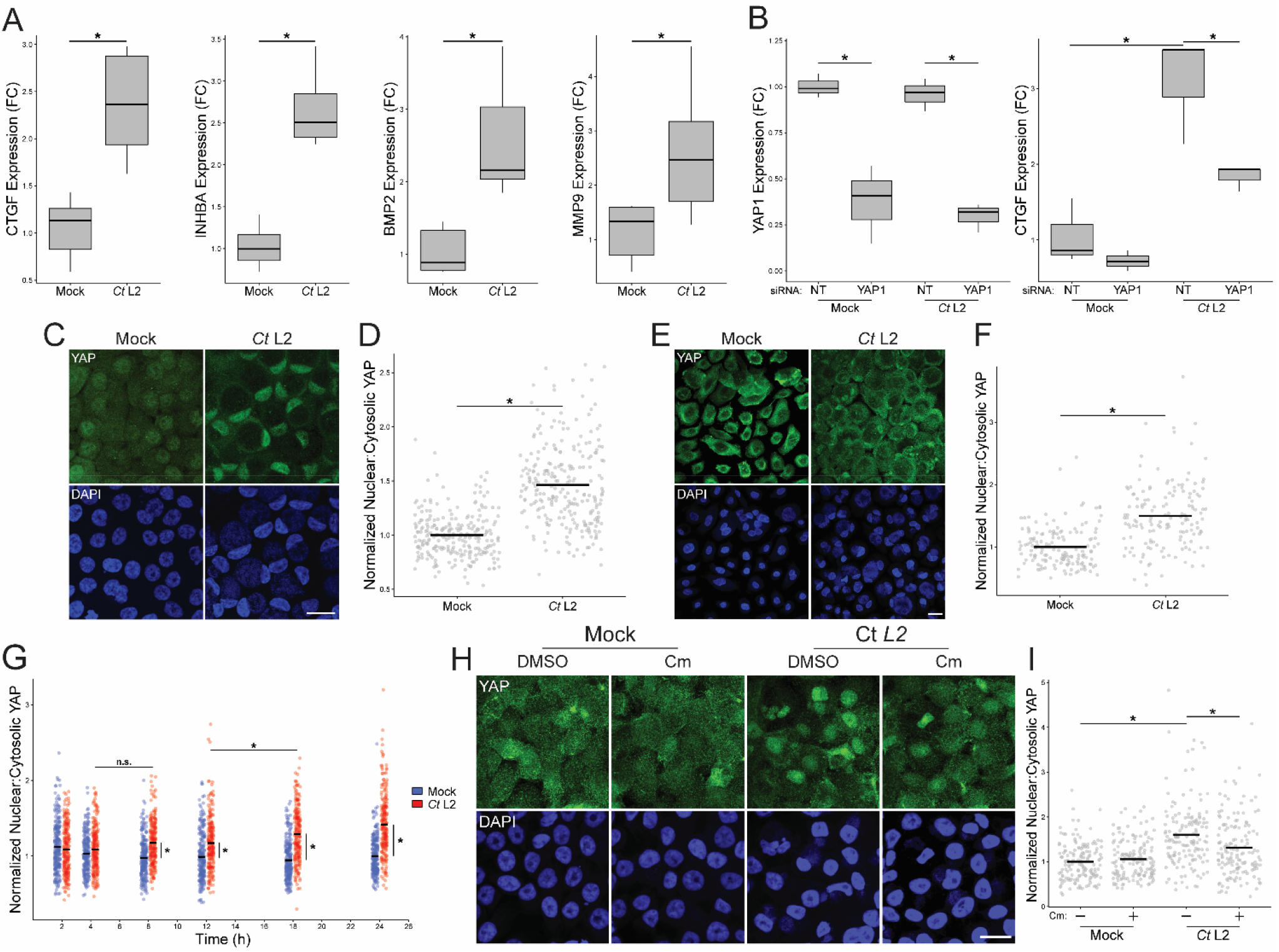
*Chlamydia* infection promotes YAP nuclear translocation. (A)Expression of CTGF, INHBA, BMP2, and and MMP9 at 24 hpi in mock- and *Ct* L2-infected End1 cells, as measured by RT-qPCR. n = 6 biological replicates; fold changes are relative to mean expression of the mock-infected and untreated control. Whiskers: minimum to maximum; asterisks: p-values ≤ 0.05, using pairwise Student’s t-tests and Bonferroni’s correction for multiple comparisons. (B)Expression of YAP1 and CTGF at 24 hpi in mock- and *Ct* L2-infected End1 cells transfected with non-targeting (NT) or YAP1-targeting siRNA (10 nM for 24 h prior to infection), as measured by RT-qPCR. n = 3 biological replicates; fold changes are relative to mean expression of the mock-infected and untreated control. Whiskers: minimum to maximum; asterisks: p-values ≤ 0.05, using pairwise Student’s t-tests and Bonferroni’s correction for multiple comparisons. (C)Representative micrographs of YAP (green) translocation into the nuclei (blue) of confluent mock- and *Ct* L2-infected End1 cells at 24 hpi. Scale bar: 20 μm. (D)Quantification of YAP nuclear translocation in (A) as a ratio of nuclear to cytosolic YAP fluorescence. n = 5 biological replicates, 50 cells measured per sample. Black bars: group means; asterisks: p-values ≤ 0.05, using pairwise Wilcoxon rank sum tests and Bonferroni’s correction for multiple comparisons. (E)Representative micrographs of YAP (green) translocation into the nuclei (blue) of confluent mock- and *Ct* L2-infected primary human cervical epithelial cells at 24 hpi. Scale bar: 20 μm. (F)Quantification of YAP nuclear translocation in (C) as a ratio of nuclear to cytosolic YAP fluorescence. n = 5 biological replicates, 50 cells measured per sample. Black bars: group means; asterisks: p-values ≤ 0.05, using pairwise Wilcoxon rank sum tests and Bonferroni’s correction for multiple comparisons. (G)Quantification of YAP nuclear translocation at 2, 4, 8, 12, 18, and 24 hpi in confluent mock- and *Ct* L2-infected End1 cells as a ratio of nuclear to cytosolic YAP fluorescence. n = 5 biological replicates, 50 cells measured per sample. Blue dots: mock-infected cells, red dots: *Ct* L2-infected cells, black bars: group means, asterisks: p-value ≤ 0.05, using pairwise Wilcoxon rank-sum tests and Bonferroni’s correction for multiple comparisons. (H)Representative micrographs of YAP (green) translocation at 18 hpi in confluent mock- and *Ct* L2-infected End1 cells treated with chloramphenicol (Cm, 50 μg/mL for 1 h at 17 hpi) or DMSO. Scale bar: 20 μm. (I)Quantification of YAP nuclear translocation in (B). n = 3 biological replicates, 50 cells measured per sample. Black bars: group means; asterisks: p-values ≤ 0.05, using pairwise Wilcoxon rank-sum tests and Bonferroni’s correction for multiple comparisons.

YAP’s ability to access the nuclear compartment and its DNA-binding partners correlates with YAP-mediated gene expression (Pocaterra et al., 2020). Thus, we opted to confirm that chlamydial infection promotes YAP activity by assaying the nuclear incidence of YAP in serovar L2-infected cells via immunofluorescence. Consistent with our initial observation of YAP target gene induction, nuclear YAP was increased in *Chlamydia-infected* End1/E6E7 cells at 24 hpi, relative to mock-infected control cells (Figure 2C). To account for variations in nuclear size and staining efficiency, we subsequently performed blind quantification of randomly selected nuclei in each sample, measuring ratios of nuclear to cytosolic YAP, analogous to previous study of YAP activation (Zhao et al., 2007; Dupont et al., 2011; Das et al., 2016). Consistently, infection was concomitant with a 1.5-fold increase in nuclear-to-cytosolic YAP ratio relative to mock-infected cells (Figure 2D). Given that expression of HPV protein E6 has been shown to stabilize YAP and thereby modulate its activity (Strickland et al., 2018), we assayed nuclear YAP translocation in primary HCECs as well, observing an equivalent increase to that in E6-expressing End1s (Figure 2E, 2F). Collectively, these data indicate chlamydial induction of YAP is irrespective of HPV E6/E7 expression.

To confirm that the observed increase in YAP activity was a *Chlamydia-driven* process requiring the action of one or more chlamydial effectors, we utilized chloramphenicol to inhibit chlamydial protein synthesis. Over the course of a 24-hour infection, we observed a significant increase in YAP nuclear localization by immunofluorescence between mock- and serovar L2-infected End1/E6E7 cells, starting as early as 8 hpi. A statistically significant increase between infected cells was first detected between 12 and 18 hpi, indicating that synthesis of YAP-modulating chlamydial effectors may occur at this stage of infection (Figure 2G). We next attempted to attenuate chlamydial YAP induction at 18 hpi via treatment with chloramphenicol. In agreement with our initial hypothesis, chloramphenicol treatment for one hour prior to fixation at 18 hpi was sufficient to significantly inhibit YAP nuclear translocation relative to a DMSO-treated control (Figures 2H, 2I). Given these data, we conclude that YAP activation in infected cells requires *de novo* synthesis of chlamydial effectors.

### 3.3 *Chlamydia* infection bypasses Hippo-mediated YAP inhibition by enhancing YAP phosphorylation at Y357

In confluent epithelial cells, YAP’s access to the nuclear compartment is inhibited by the Hippo kinase complex, which in turn is stabilized by cell-cell contact (Gumbiner and Kim, 2014). The phosphorylation cascade of Hippo complex members terminates in YAP phosphorylation at S127 and other serine residues, which drives cytoplasmic sequestration of YAP via binding to 14-3-3 (Piccolo et al., 2014). To determine if *Chlamydia* induced YAP activity by relieving Hippo-mediated inhibition, we examined YAP activation in experimentally enforced culture conditions of inhibited cell-cell contact. By seeding End1/E6E7 cells on glass treated with fibronectin in 30- and 45-μm micropatterns (Degot et al., 2010), the resulting deposition of small cell clusters allowed us to assay YAP nuclear translocation in an environment where cell-cell contact (and, by extension, Hippo-mediated YAP inhibition) was absent or severely limited. In both 30-μm patterns (1-3 cells/cluster) and 45-μm patterns (2-4 cells/cluster), we observed a significant increase in YAP nuclear translocation in serovar L2-infected cells at 24 hpi, relative to mock-infected controls (Figure 3A). Quantification of nuclear translocation by immunofluorescence indicated that infection promotes a 1.5-fold increase in YAP activation relative to uninfected cells, equivalent to our observations in confluent monolayers (Figure 3B). This phenotype was additionally observed when adherens junctions were disassembled by calcium chloride withdrawal in confluent *Chlamydia-infected* End1/E6E7 monolayers (Supplementary Figure S2) (O’Keefe et al., 1987). Taken together, these data suggest that chlamydial activation of YAP occurs independent of cell-cell contact, indicating this phenotype is not dependent on destabilization of the YAP-inhibitory Hippo kinase complex.

**Figure 3:**
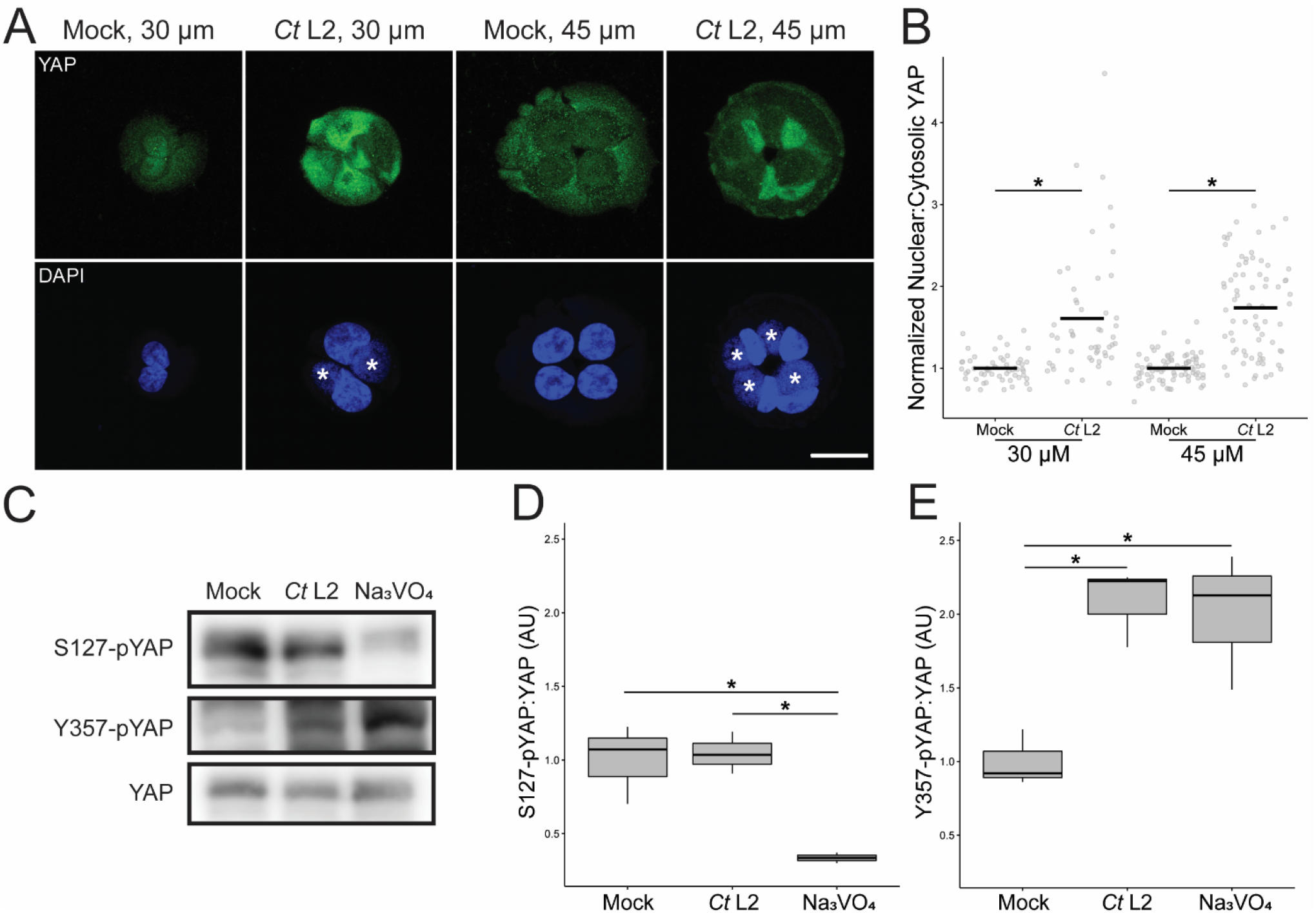
*Chlamydia* infection bypasses Hippo-mediated YAP inhibition by enhancing YAP phosphorylation at Y357. (A)Representative micrographs of YAP (green) translocation into the nuclei (blue) at 24 hpi of mock- and *Ct* L2-infected End1 cells cultured in small cell clusters using 30 μm (1-2 cells) and 45 μm (2-4 cells) micropatterns. Asterisks: chlamydial inclusions; scale bar: 20 μm. (B)Quantification of YAP nuclear translocation in (A) as a ratio of nuclear to cytosolic YAP fluorescence. n = 3 biological replicates, with a minimum of 5 fields measured per sample. Black bars: group means; asterisks: p-values ≤ 0.05, using pairwise Wilcoxon rank sum tests and Bonferroni’s correction for multiple comparisons. (C)Representative Western blot of YAP phosphorylation of mock- and *Ct* L2-infected or sodium orthovanadate-treated (Na_3_VO_4_, 10 mM for 1 h at 23 hpi) End1 cells at 24 hpi. (D, E) Densitometric quantification of YAP phosphorylation at S127 (D) and Y357 (E), normalized to total YAP. n = 3 biological replicates; whiskers: minimum to maximum; asterisks: p-values ≤ 0.05, using pairwise Student’s t-tests and Bonferroni’s correction for multiple comparisons.

The recruitment of Hippo cascade members to adherens junctions promotes the formation of a YAP-inhibitory complex, resulting in YAP phosphorylation at serine residues by Hippo complex members LATS1/2 (Zhao et al., 2007). In contrast, recent studies have implicated phosphorylation of YAP at Y357 and other tyrosine residues in Hippo-independent YAP activation (Elbediwy et al., 2018; Sugihara et al., 2018, 2018). To confirm that chlamydial YAP activation occurs via Hippoin-dependent signaling, we thus surveyed YAP S127 and Y357 phosphorylation in confluent, *Chlamydia-infected* End1s via Western blotting. Serovar L2-infected End1/E6E7 cells exhibited increased YAP Y357 phosphorylation at 24 hpi relative to a mock-infected control. This result was consistent with mock-infected cells treated with sodium orthovanadate (Na_3_VO_4_), a tyrosine phosphatase inhibitor that serves as a positive control for increased YAP tyrosine phosphorylation (Figure 3C) (Sugihara et al., 2018). Surprisingly, L2-infected cells did not exhibit altered YAP S127, suggesting infection does not relieve Hippo-mediated YAP inhibition (Figure 3D). By contrast, pY357-YAP levels were increased in an infection-dependent manner (Figure 3E). Collectively, these data demonstrate that infection-dependent YAP activation bypasses S127 phospho-inhibition by the Hippo kinase cascade, instead promoting alternative phospho-activation of YAP at Y357 to facilitate its nuclear translocation.

### 3.4 SFK-mediated YAP phosphorylation at Y357 is required for chlamydial YAP activation

In contrast with the YAP-inhibitory effect of Hippo kinase at cell-cell junctions, YAP activation has been associated with cell-substrate contact, as well as the activity of Src-family kinases. Cell adhesion to fibronectin has been shown to promote YAP nuclear translocation in a Src-dependent, Hippo-inhibitory fashion (Kim and Gumbiner, 2015). Intriguingly, Src and the related kinase Lck are also implicated in Hippo-independent YAP activation via Y357 phosphorylation (Li et al., 2016; Elbediwy et al., 2018; Smoot et al., 2018; Sugihara et al., 2018). Further, it has been shown that *Chlamydia* induces Src activity (Mital and Hackstadt, 2011), and that Src-family kinase (SFK) modulation is critical to multiple stages of chlamydial pathogenesis (Elwell et al., 2011). Thus, to determine if infection-mediated YAP activity was the product of SFK-mediated YAP activation, we assayed YAP nuclear translocation of infected cells treated with the pan-SFK inhibitor PP2. Consistent with our hypothesis, serovar L2-infected cells treated with PP2 over the course of a 24 hour infection exhibited statistically significant attenuation of YAP nuclear translocation relative to a DMSO-treated control (Figures 4A, 4B). YAP phosphorylation at Y357 in infected cells was also severely attenuated by SFK inhibition (Figure 4C), consistent with studies directly implicating SFKs in YAP tyrosine phosphorylation (Li et al., 2016; Sugihara et al., 2018). To confirm that YAP phosphorylation at this residue is indeed required for infection-dependent YAP nuclear translocation, we ectopically expressed FLAG-tagged, Y357F-mutant YAP in infected End1s, comparing nuclear translocation to an equivalent mock-infected control. Importantly, nuclear translocation of the phospho-dead YAP-Y357F mutant in infected cells was indistinguishable from mock-infected cells expressing the same construct, whereas infection significantly increased the nuclear translocation of FLAG-tagged, wild-type YAP relative to a mock-infected control (Figures 4D, 4E). Taken together, these data indicate that chlamydial induction of YAP activity operates via SFK- and Y357-dependent means of enhancing YAP nuclear translocation.

**Figure 4:**
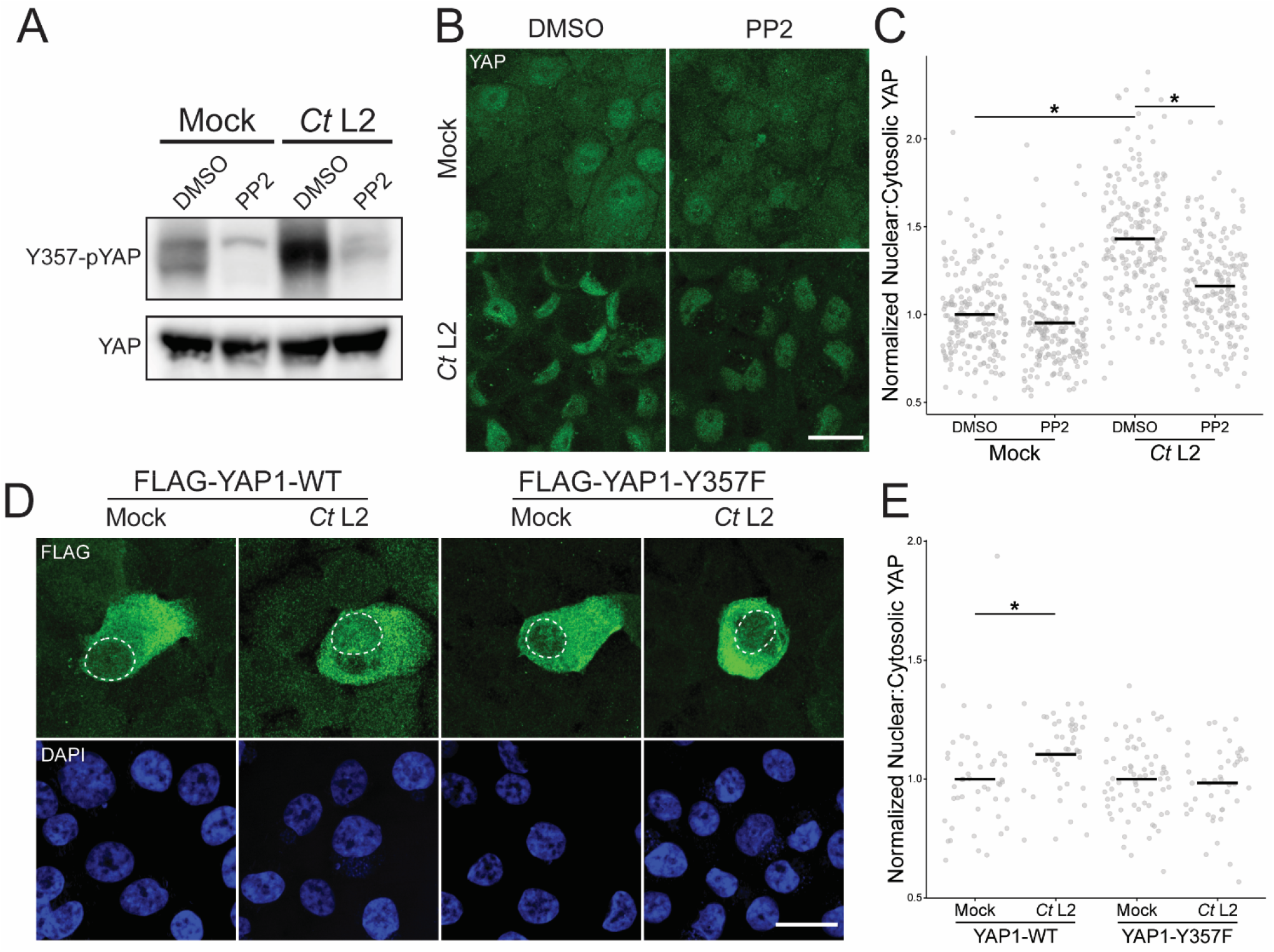
Infection-induced phosphorylation at Y357 is required for YAP nuclear translocation. (A)Representative Western blot of YAP tyrosine phosphorylation of mock- and *Ct* L2-infected End1 cells at 24 hpi, treated with the Src-family kinase inhibitor PP2 (10 μM for 24 h), or DMSO. (B)Representative micrographs of YAP translocation at 24 hpi in confluent mock- and *Ct* L2-infected End1 cells treated with the Src-family kinase inhibitor PP2 (10 μM for 24 h), or DMSO. Scale bar: 20 μm. (C)Quantification of YAP nuclear translocation in (B). n = 4 biological replicates, 50 cells measured per sample. Black bars: group means; asterisks: p-values ≤ 0.05, using pairwise Wilcoxon rank sum tests and Bonferroni’s correction for multiple comparisons. (D)Representative micrographs of the translocation of FLAG-tagged YAP wild-type or the phospho-dead Y357F mutant (green) into the nuclei (blue) of confluent mock- and *Ct* L2-infected End1 cells at 24 hpi. Scale bar: 20 μm, dotted lines: nuclear area of transfected cells. (E)Quantification of YAP nuclear translocation in (D). n = 4 biological replicates, 50 cells measured per sample. Black bars: group means; asterisks: p-values ≤ 0.05, using pairwise Wilcoxon rank sum tests and Bonferroni’s correction for multiple comparisons.

## 4 Discussion

In this report, we present a novel mechanism for *Chlamydia*-directed modulation of host gene expression. Infection of primary and immortalized epithelial cells with *C. trachomatis* serovar L2 resulted in a host transcriptome consistent with induction of the transcriptional coactivator YAP and its DNA-binding partners. Consistent with this observation, infected cells also exhibit increased YAP nuclear translocation. This phenotype was concomitant with increased expression of YAP target genes INHBA, BMP2, MMP9, and CTGF (which in the latter case was sensitive to YAP siRNA-mediated knockdown). Enhanced YAP activation is first detectable at 18 hours post-infection; critically, this phenotype was attenuated by inhibition of bacterial protein synthesis, indicating YAP activation is a pathogen-directed effect of infection. Intriguingly, *Chlamydia* infection induced YAP nuclear translocation in epithelial cells that were sparsely seeded on micropatterned chips – a scenario in which canonical YAP inhibition by the Hippo kinase cascade is minimal – suggesting that infection enhances YAP activation in a Hippo-independent fashion. Accordingly, Hippo-dependent YAP phospho-inhibition at S127 was unaffected by infection, whereas YAP phosphorylation at Y357 was enhanced. Inhibition of host Src-family kinases significantly attenuated both YAP Y357 phosphorylation and nuclear translocation, consistent with previous reports of SFK-mediated phosphorylation at this residue enhancing YAP activity. Importantly, we observe that infection does not promote nuclear translocation of a YAP Y357F mutant, indicating that phosphorylation at this residue is required for chlamydial YAP activation. Taken together, our data show *Chlamydia* induces YAP-dependent gene expression by acting upon an alternative, Hippo-independent regulatory pathway in the host, involving phosphorylation of YAP at Y357 and the activity of host Src-family kinases.

Our observation of *Chlamydia*-directed induction of YAP activity is consistent with prior reports of YAP modulation by other bacterial pathogens. The Gram-negative pathogen *Helicobacter pylori* was recently shown to enhance YAP expression through the action of the effector CagA (Li et al., 2018). It is unclear how CagA interacts with the steady-state inhibition of YAP by the Hippo kinase cascade; given the prior finding that CagA destabilizes host cell-cell junctions – a common site of Hippo kinase complex recruitment – one possible explanation is that CagA antagonizes Hippo-mediated YAP inhibition by indirectly preventing stabilization of the Hippo complex (Song et al., 2013). Alternatively, the observation that YAP expression is increased by CagA in a dose-dependent fashion suggests that *H. pylori* infection may instead circumvent Hippo inhibition by overwhelming either Hippo’s capacity to phospho-inhibit YAP or 14-3-3’s capacity to sequester serine-phosphorylated YAP in the cytoplasm (Li et al., 2018). The observation that *H. pylori* enhances EMT in gastric epithelia in a CagA-dependent fashion suggests a potential for pathogen-directed YAP activation to have downstream effects on pathology (Song et al., 2013; Li et al., 2018). Regardless, these results illustrate that a component of the altered host transcriptome observed during infection is the product of pathogen manipulation of host transcription factors, rather than a general host response to the pathogen.

Our finding that chlamydial YAP activation is sensitive to inhibition of bacterial protein synthesis indicates that one or more chlamydial effectors facilitate YAP activation. One possible candidate for this effect is the chlamydial inclusion protein IncG, which has been previously shown to interact with host 14-3-3β (Scidmore and Hackstadt, 2001). Given that 14-3-3-family proteins are known to act downstream of Hippo-mediated YAP inhibition by binding and sequestering serine-phosphorylated YAP in the cytosol (Basu et al., 2003; Zhao et al., 2007), potential inhibition of 14-3-3 by IncG may complement infection-dependent Y357 phosphorylation in decoupling YAP from Hippo-mediated phospho-inhibition at S127. Additional YAP-activating factors may operate via enhancement of YAP tyrosine phosphorylation, an event observed to be essential for infection-mediated YAP translocation. Given that chlamydial YAP phospho-activation is demonstrably sensitive to inhibition of Src-family kinases, screening for chlamydial modulators of SFKs or signaling pathways that lead to increased SFK activity may identify additional YAP-activating effectors.

Although we do not observe chlamydial inhibition of the Hippo kinase cascade at the level of YAP S127 phosphorylation, we cannot entirely dismiss a role for this negative regulator in determining levels of nuclear YAP during infection. The recent report of chlamydial disruption of cell-cell junctions in an organoid model of infection (Dolat and Valdivia, 2021) suggests that YAP inhibition by Hippo may be relevant in three-dimensional models of infection. Our laboratory has previously reported on chlamydial modulation of focal adhesion kinase (FAK) (Thwaites et al., 2014; Pedrosa et al., 2020); given that FAK has been shown to antagonize Hippo-mediated YAP phosphorylation at S397 (Hu et al., 2017), infection may potentially impact YAP inhibition by Hippo in this way as well. Though beyond the scope of this initial study, further investigation of chlamydial YAP activation will likely require use of three-dimensional culture models to determine which of the extensive portfolio of subcellular structures and mechanical inputs co-opted by the pathogen ultimately drive enhancement of YAP activity during infection.

The effect of *Chlamydia-induced* YAP nuclear translocation on host gene expression is still somewhat unclear. Critically, a substantial component of the YAP-responsive gene set was downregulated in response to infection – an outcome inconsistent with YAP’s reported function as a transcriptional coactivator. One explanation for this result is the activity of transcriptional repressors induced by infection: while infection-downregulated genes designated by ChEA as YAP targets may be regulated YAP-cofactor complexes, this event may be superseded by infection-induced repressors also binding to the same promoters. Ultimately, future work to define the true extent of the *Chlamydia*-induced YAP regulon will require comparing the infected host transcriptome reported here to an equivalent infection in a YAP-negative background. Given that the relatively modest siRNA-mediated knockdown we report here is consistent with past use of siRNA in End1s (Govender et al., 2014), generating a CRISPR/Cas9-mediated YAP-knockout from an alternative cell line may be necessary.

Our results illustrate a mechanism by which chlamydial infection promotes YAP-dependent gene expression in the host; however, the benefit of this interaction to the pathogen is unclear. YAP activation in infected epithelial cells can be viewed as a downstream process relevant to pathogenesis, but not necessarily one beneficial to chlamydial growth in host cells. Given past work demonstrating *Chlamydia-infected* cells exhibit increased expression of genes encoding pro-fibrotic cytokines and ECM components (Humphrys et al., 2013; Porcella et al., 2015), and our own observation that infection induces expression of fibroblast-activating signaling factors (CTGF, INHBA, BMP2), the potential effect of infection-mediated YAP activation on *Chlamydia*-associated scarring warrants further study. Fibroblasts in particular are understood to play a critical part in other types of scar-forming disease (Kendall and Feghali-Bostwick, 2014); however, their contribution to chlamydial fibrosis has yet to be fully assessed. Understanding the yet uncharacterized effect of *Chlamydia*-infected cells on fibroblasts and other uninfected cell types will likely be critical to elucidating mechanisms of chlamydial fibrosis *in vivo*, as well as contextualizing the broader role of YAP induction in chlamydial pathogenesis.

## 5 Acknowledgments

The authors acknowledge the members of the Carabeo laboratory for their critical feedback on the development of this project. We also acknowledge the generosity of Dr. Ted Hackstadt for providing the chlamydial strain used in this study, Dr. Marius Sudol for providing pEGFP-C3-HYAP1, as well as the WSU SMB Genomics Core and UNMC Genomics Core for performing mRNA library preparation and bulk RNA-sequencing, and the UNMC Tissue Science Facility for RAFT culture sectioning and slide preparation.

## 6 Contribution

Our study presents a means by which *C. trachomatis* induces gene expression in host epithelial cells, providing an explanation for how infection mediates a host transcriptomic response above and beyond pathogen recognition. We identify chlamydial induction of YAP activity as a principal mediator of this effect, and thus a potential therapeutic target for *Chlamydia*-associated pathology. Our finding that infection bypasses Hippo-mediated YAP inhibition by enhancing YAP Y357 phosphorylation in a pathogen-directed, Src-family kinase-dependent fashion corroborates recent discoveries indicating this relatively novel axis of YAP activation can circumvent its canonical inhibition by the Hippo kinase cascade. Collectively, our data illustrates an underappreciated aspect of the host-pathogen interaction: pathogen-directed regulation of transcription factor activity driving changes in the gene expression of the host.

## 7 Conflict of Interest

The authors declare that the research was conducted in the absence of any commercial or financial relationships that could be construed as a potential conflict of interest.

## 8 Author Contributions

Conceptualization, L.C. and R.C.; Methodology, L.C., A.B., and R.C.; Validation, L.C. and A.B.; Formal Analysis, L.C.; Investigation, L.C. and A.B.; Resources, R.C.; Data Curation, L.C.; Writing – Original Draft, L.C.; Writing – Review & Editing, R.C. and A.B.; Visualization, L.C.; Supervision, R.C.; Project Administration, R.C.; Funding Acquisition, R.C.

## 9 Funding

This work was supported by NIAID grant R01 AI065545 to RAC. The University of Nebraska DNA Sequencing Core receives partial support from the National Institute for General Medical Science (NIGMS) INBRE - P20GM103427-19 grant as well as The Fred & Pamela Buffett Cancer Center Support Grant - P30 CA036727. This publication’s contents are the sole responsibility of the authors, and do not necessarily represent the official views of the NIH.

## 11 Data Availability Statement

The bulk RNA-sequencing datasets generated and analyzed for this study can be found in the Gene Expression Omnibus (Edgar et al., 2002), GEO Series accession number GSE180784 (https://www.ncbi.nlm.nih.gov/geo/query/acc.cgi?acc=GSE180784).

## Notes

https://www.ncbi.nlm.nih.gov/geo/query/acc.cgi?acc=GSE180784

### Competing Interest Statement

The authors have declared no competing interest.

### Summary of Updates

Reduction in scope to focus exclusively on the effect of infection on YAP activity; Section on infection-mediated YAP target gene induction updated to include expression data from infection primary epithelial cells; Section on infection promoting YAP nuclear translocation updated to include complementary data in primary cells; Section on infection-mediated YAP activation requiring Y357 phosphorylation updated to include nuclear translocation of Y357F mutant YAP; general formatting updates for submission to Frontiers.

## References

Aasen, T., Hodgins, M. B., Edward, M., and Graham, S. V. (2003). The relationship between connexins, gap junctions, tissue architecture and tumour invasion, as studied in a novel in vitro model of HPV-16-associated cervical cancer progression. Oncogene 22, 7969–7980. doi: 10.1038/sj.onc.1206709.

Basu, S., Totty, N. F., Irwin, M. S., Sudol, M., and Downward, J. (2003). Akt phosphorylates the Yes-associated protein, YAP, to induce interaction with 14-3-3 and attenuation of p73-mediated apoptosis. Mol Cell 11, 11–23. doi: 10.1016/s1097-2765(02)00776-1.

Caldwell, H. D., Kromhout, J., and Schachter, J. (1981). Purification and partial characterization of the major outer membrane protein of Chlamydia trachomatis. Infect Immun 31, 1161–1176.

Carlson, J. H., Porcella, S. F., McClarty, G., and Caldwell, H. D. (2005). Comparative genomic analysis of Chlamydia trachomatis oculotropic and genitotropic strains. Infect. Immun. 73, 6407–6418. doi: 10.1128/IAI.73.10.6407-6418.2005.

Cheng, W., Shivshankar, P., Zhong, Y., Chen, D., Li, Z., and Zhong, G. (2008). Intracellular Interleukin-1α Mediates Interleukin-8 Production Induced by Chlamydia trachomatis Infection via a Mechanism Independent of Type I Interleukin-1 Receptor. Infect. Immun. 76, 942–951. doi: 10.1128/IAI.01313-07.

Chow, J. M., Yonekura, M. L., Richwald, G. A., Greenland, S., Sweet, R. L., and Schachter, J. (1990). The association between Chlamydia trachomatis and ectopic pregnancy. A matched-pair, case-control study. JAMA 263, 3164–3167.

Christian, J., Vier, J., Paschen, S. A., and Häcker, G. (2010). Cleavage of the NF-κB Family Protein p65/RelA by the Chlamydial Protease-like Activity Factor (CPAF) Impairs Proinflammatory Signaling in Cells Infected with Chlamydiae *. Journal of Biological Chemistry 285, 41320–41327. doi: 10.1074/jbc.M110.152280.

Das, A., Fischer, R. S., Pan, D., and Waterman, C. M. (2016). YAP Nuclear Localization in the Absence of Cell-Cell Contact Is Mediated by a Filamentous Actin-dependent, Myosin II- and Phospho-YAP-independent Pathway during Extracellular Matrix Mechanosensing. J Biol Chem 291, 6096–6110. doi: 10.1074/jbc.M115.708313.

Degot, S., Auzan, M., Chapuis, V., Béghin, A., Chadeyras, A., Nelep, C., et al. (2010). Improved Visualization and Quantitative Analysis of Drug Effects Using Micropatterned Cells. J Vis Exp, 2514. doi: 10.3791/2514.

Dolat, L., and Valdivia, R. H. (2021). An endometrial organoid model of interactions between Chlamydia and epithelial and immune cells. Journal of Cell Science 134, jcs252403. doi: 10.1242/jcs.252403.

Dupont, S., Morsut, L., Aragona, M., Enzo, E., Giulitti, S., Cordenonsi, M., et al. (2011). Role of YAP/TAZ in mechanotransduction. Nature 474, 179–183. doi: 10.1038/nature10137.

Eckmann, L., Kagnoff, M. F., and Fierer, J. (1993). Epithelial cells secrete the chemokine interleukin-8 in response to bacterial entry. Infect Immun 61, 4569–4574. doi: 10.1128/iai.61.11.4569-4574.1993.

Edgar, R., Domrachev, M., and Lash, A. E. (2002). Gene Expression Omnibus: NCBI gene expression and hybridization array data repository. Nucleic Acids Res 30, 207–210.

Elbediwy, A., Vanyai, H., Diaz-de-la-Loza, M.-C., Frith, D., Snijders, A. P., and Thompson, B. J. (2018). Enigma proteins regulate YAP mechanotransduction. J Cell Sci 131, jcs221788. doi: 10.1242/jcs.221788.

Elwell, C. A., Kierbel, A., and Engel, J. N. (2011). Species-Specific Interactions of Src Family Tyrosine Kinases Regulate Chlamydia Intracellular Growth and Trafficking. mBio 2, e00082–11. doi: 10.1128/mBio.00082-11.

Fichorova, R. N., Rheinwald, J. G., and Anderson, D. J. (1997). Generation of Papillomavirus-Immortalized Cell Lines from Normal Human Ectocervical, Endocervical, and Vaginal Epithelium that Maintain Expression of Tissue-Specific Differentiation Proteins. Biol Reprod 57, 847–855. doi: 10.1095/biolreprod57.4.847.

Futakuchi, A., Inoue, T., Wei, F.-Y., Inoue-Mochita, M., Fujimoto, T., Tomizawa, K., et al. (2018). YAP/TAZ Are Essential for TGF-β2-Mediated Conjunctival Fibrosis. Invest Ophthalmol Vis Sci 59, 3069–3078. doi: 10.1167/iovs.18-24258.

Gali, Y., Ariën, K. K., Praet, M., Van den Bergh, R., Temmerman, M., Delezay, O., et al. (2010). Development of an in vitro dual-chamber model of the female genital tract as a screening tool for epithelial toxicity. Journal of Virological Methods 165, 186–197. doi: 10.1016/j.jviromet.2010.01.018.

Govender, Y., Avenant, C., Verhoog, N. J. D., Ray, R. M., Grantham, N. J., Africander, D., et al. (2014). The injectable-only contraceptive medroxyprogesterone acetate, unlike norethisterone acetate and progesterone, regulates inflammatory genes in endocervical cells via the glucocorticoid receptor. PLoS One 9, e96497. doi: 10.1371/journal.pone.0096497.

Gumbiner, B. M., and Kim, N.-G. (2014). The Hippo-YAP signaling pathway and contact inhibition of growth. J Cell Sci 127, 709–717. doi: 10.1242/jcs.140103.

Haggerty, C. L., Gottlieb, S. L., Taylor, B. D., Low, N., Xu, F., and Ness, R. B. (2010a). Risk of Sequelae after *Chlamydia trachomatis* Genital Infection in Women. The Journal of Infectious Diseases 201, 134–155. doi: 10.1086/652395.

Haggerty, C. L., Gottlieb, S. L., Taylor, B. D., Low, N., Xu, F., and Ness, R. B. (2010b). Risk of Sequelae after Chlamydia trachomatis Genital Infection in Women. The Journal of Infectious Diseases 201, S134–S155. doi: 10.1086/652395.

Hu, J. K.-H., Du, W., Shelton, S. J., Oldham, M. C., DiPersio, C. M., and Klein, O. D. (2017). A FAK-YAP-mTOR signaling axis regulates stem cell-based tissue renewal in mice. Cell Stem Cell 21, 91–106.e6. doi: 10.1016/j.stem.2017.03.023.

Hu, V. H., Harding-Esch, E. M., Burton, M. J., Bailey, R. L., Kadimpeul, J., and Mabey, D. C. W. (2010). Epidemiology and control of trachoma: systematic review. Trop Med Int Health 15, 673–691. doi: 10.1111/j.1365-3156.2010.02521.x.

Huang, Z., Hu, J., Pan, J., Wang, Y., Hu, G., Zhou, J., et al. (2016). YAP stabilizes SMAD1 and promotes BMP2-induced neocortical astrocytic differentiation. Development 143, 2398–2409. doi: 10.1242/dev.130658.

Humphrys, M. S., Creasy, T., Sun, Y., Shetty, A. C., Chibucos, M. C., Drabek, E. F., et al. (2013). Simultaneous Transcriptional Profiling of Bacteria and Their Host Cells. PLOS ONE 8, e80597. doi: 10.1371/journal.pone.0080597.

Johnson, K. A., Lee, J. K., Chen, A. L., Tan, M., and Sütterlin, C. (2015). Induction and inhibition of CPAF activity during analysis of Chlamydia-infected cells. Pathog Dis 73, 1–8. doi: 10.1093/femspd/ftv007.

Kendall, R. T., and Feghali-Bostwick, C. A. (2014). Fibroblasts in fibrosis: novel roles and mediators. Frontiers in Pharmacology 5. Available at: https://www.frontiersin.org/article/10.3389/fphar.2014.00123 [Accessed March 15, 2022].

Kim, N.-G., and Gumbiner, B. M. (2015). Adhesion to fibronectin regulates Hippo signaling via the FAK-Src-PI3K pathway. J Cell Biol 210, 503–515. doi: 10.1083/jcb.201501025.

Kiviat, N. B., Wølner-Hanssen, P., Eschenbach, D. A., Wasserheit, J. N., Paavonen, J. A., Bell, T. A., et al. (1990). Endometrial histopathology in patients with culture-proved upper genital tract infection and laparoscopically diagnosed acute salpingitis. Am. J. Surg. Pathol. 14, 167–175.

Krämer, S., Crauwels, P., Bohn, R., Radzimski, C., Szaszák, M., Klinger, M., et al. (2015). AP-1 Transcription Factor Serves as a Molecular Switch between Chlamydia pneumoniae Replication and Persistence. Infect Immun 83, 2651–2660. doi: 10.1128/IAI.03083-14.

Lachmann, A., Xu, H., Krishnan, J., Berger, S. I., Mazloom, A. R., and Ma’ayan, A. (2010). ChEA: transcription factor regulation inferred from integrating genome-wide ChIP-X experiments. Bioinformatics 26, 2438–2444. doi: 10.1093/bioinformatics/btq466.

Lad, S. P., Fukuda, E. Y., Li, J., de la Maza, L. M., and Li, E. (2005). Up-regulation of the JAK/STAT1 signal pathway during Chlamydia trachomatis infection. J Immunol 174, 7186–7193. doi: 10.4049/jimmunol.174.11.7186.

Lansingh, V. C. (2016). Trachoma. BMJ Clin Evid 2016, 0706.

Le Negrate, G., Krieg, A., Faustin, B., Loeffler, M., Godzik, A., Krajewski, S., et al. (2008). ChlaDub1 of Chlamydia trachomatis suppresses NF-κB activation and inhibits IκBα ubiquitination and degradation. Cellular Microbiology 10, 1879–1892. doi: 10.1111/j.1462-5822.2008.01178.x.

Lee, J. K., Enciso, G. A., Boassa, D., Chander, C. N., Lou, T. H., Pairawan, S. S., et al. (2018). Replication-dependent size reduction precedes differentiation in Chlamydia trachomatis. Nat Commun 9, 45. doi: 10.1038/s41467-017-02432-0.

Li, C., Wen, A., Shen, B., Lu, J., Huang, Y., and Chang, Y. (2011). FastCloning: a highly simplified, purification-free, sequence- and ligation-independent PCR cloning method. BMC Biotechnology 11, 92. doi: 10.1186/1472-6750-11-92.

Li, N., Feng, Y., Hu, Y., He, C., Xie, C., Ouyang, Y., et al. (2018). Helicobacter pylori CagA promotes epithelial mesenchymal transition in gastric carcinogenesis via triggering oncogenic YAP pathway. J Exp Clin Cancer Res 37, 280. doi: 10.1186/s13046-018-0962-5.

Li, P., Silvis, M. R., Honaker, Y., Lien, W.-H., Arron, S. T., and Vasioukhin, V. (2016). αE-catenin inhibits a Src-YAP1 oncogenic module that couples tyrosine kinases and the effector of Hippo signaling pathway. Genes Dev 30, 798–811. doi: 10.1101/gad.274951.115.

Mital, J., and Hackstadt, T. (2011). Diverse Requirements for Src-Family Tyrosine Kinases Distinguish Chlamydial Species. mBio. doi: 10.1128/mBio.00031-11.

Mo, J.-S., Yu, F.-X., Gong, R., Brown, J. H., and Guan, K.-L. (2012). Regulation of the Hippo-YAP pathway by protease-activated receptors (PARs). Genes Dev 26, 2138–2143. doi: 10.1101/gad.197582.112.

Ness, R. B., Soper, D. E., Richter, H. E., Randall, H., Peipert, J. F., Nelson, D. B., et al. (2008). Chlamydia Antibodies, Chlamydia Heat Shock Protein, and Adverse Sequelae After Pelvic Inflammatory Disease: The PID Evaluation and Clinical Health (PEACH) Study. Sexually Transmitted Diseases 35, 129–135. doi: 10.1097/OLQ.0b013e3181557c25.

Nogueira, A. T., Braun, K. M., and Carabeo, R. A. (2017). Characterization of the Growth of Chlamydia trachomatis in In Vitro-Generated Stratified Epithelium. Front Cell Infect Microbiol 7, 438. doi: 10.3389/fcimb.2017.00438.

O’Keefe, E. J., Briggaman, R. A., and Herman, B. (1987). Calcium-induced assembly of adherens junctions in keratinocytes. J Cell Biol 105, 807–817. doi: 10.1083/jcb.105.2.807.

Pan, Z., Tian, Y., Zhang, B., Zhang, X., Shi, H., Liang, Z., et al. (2017). YAP signaling in gastric cancer-derived mesenchymal stem cells is critical for its promoting role in cancer progression. Int J Oncol 51, 1055–1066. doi: 10.3892/ijo.2017.4101.

Pedrosa, A. T., Murphy, K. N., Nogueira, A. T., Brinkworth, A. J., Thwaites, T. R., Aaron, J., et al. (2020). A post-invasion role for Chlamydia type III effector TarP in modulating the dynamics and organization of host cell focal adhesions. J Biol Chem 295, 14763–14779. doi: 10.1074/jbc.RA120.015219.

Piccolo, S., Dupont, S., and Cordenonsi, M. (2014). The biology of YAP/TAZ: hippo signaling and beyond. Physiol Rev 94, 1287–1312. doi: 10.1152/physrev.00005.2014.

Pocaterra, A., Romani, P., and Dupont, S. (2020). YAP/TAZ functions and their regulation at a glance. Journal of Cell Science 133. doi: 10.1242/jcs.230425.

Porcella, S. F., Carlson, J. H., Sturdevant, D. E., Sturdevant, G. L., Kanakabandi, K., Virtaneva, K., et al. (2015). Transcriptional Profiling of Human Epithelial Cells Infected with Plasmid Bearing and Plasmid-Deficient Chlamydia trachomatis. Infect. Immun. 83, 534–543. doi: 10.1128/IAI.02764-14.

Price, M. J., Ades, A. E., Soldan, K., Welton, N. J., Macleod, J., Simms, I., et al. (2016). Pelvic inflammatory disease and tubal factor infertility. NIHR Journals Library Available at: https://www.ncbi.nlm.nih.gov/books/NBK350656/ [Accessed April 7, 2018].

Rasmussen, S. J., Eckmann, L., Quayle, A. J., Shen, L., Zhang, Y. X., Anderson, D. J., et al. (1997). Secretion of proinflammatory cytokines by epithelial cells in response to Chlamydia infection suggests a central role for epithelial cells in chlamydial pathogenesis. J Clin Invest 99, 77–87.

Scidmore, M. A., and Hackstadt, T. (2001). Mammalian 14-3-3beta associates with the Chlamydia trachomatis inclusion membrane via its interaction with IncG. Mol Microbiol 39, 1638–1650. doi: 10.1046/j.1365-2958.2001.02355.x.

Smoot, R. L., Werneburg, N. W., Sugihara, T., Hernandez, M. C., Yang, L., Mehner, C., et al. (2018). Platelet-derived growth factor regulates YAP transcriptional activity via Src family kinase dependent tyrosine phosphorylation. Journal of Cellular Biochemistry 119, 824–836. doi: https://doi.org/10.1002/jcb.26246.

Snavely, E. A., Kokes, M., Dunn, J. D., Saka, H. A., Nguyen, B. D., Bastidas, R. J., et al. (2014). Reassessing the role of the secreted protease CPAF in Chlamydia trachomatis infection through genetic approaches. Pathog Dis 71, 336–351. doi: 10.1111/2049-632X.12179.

Song, X., Chen, H.-X., Wang, X.-Y., Deng, X.-Y., Xi, Y.-X., He, Q., et al. (2013). H. pylori-encoded CagA disrupts tight junctions and induces invasiveness of AGS gastric carcinoma cells via Cdx2-dependent targeting of Claudin-2. Cell Immunol 286, 22–30. doi: 10.1016/j.cellimm.2013.10.008.

Stephens, R. S. (2003). The cellular paradigm of chlamydial pathogenesis. Trends in Microbiology 11, 44–51. doi: 10.1016/S0966-842X(02)00011-2.

Strickland, S., Brimer, N., Lyons, C., and Vande Pol, S. B. (2018). Human Papillomavirus E6 interaction with cellular PDZ domain proteins modulates YAP nuclear localization. Virology 516, 127–138. doi: 10.1016/j.virol.2018.01.003.

Sugihara, T., Werneburg, N. W., Hernandez, M. C., Yang, L., Kabashima, A., Hirsova, P., et al. (2018). YAP Tyrosine Phosphorylation and Nuclear Localization in Cholangiocarcinoma Cells Are Regulated by LCK and Independent of LATS Activity. Mol Cancer Res 16, 1556–1567. doi: 10.1158/1541-7786.MCR-18-0158.

Sun, Y., Zhou, P., Chen, S., Hu, C., Bai, Q., Wu, H., et al. (2017). The JAK/STAT3 signaling pathway mediates inhibition of host cell apoptosis by Chlamydia psittaci infection. Pathog Dis 75. doi: 10.1093/femspd/ftx088.

Szeto, S. G., Narimatsu, M., Lu, M., He, X., Sidiqi, A. M., Tolosa, M. F., et al. (2016). YAP/TAZ Are Mechanoregulators of TGF-β-Smad Signaling and Renal Fibrogenesis. JASN, ASN.2015050499. doi: 10.1681/ASN.2015050499.

Taylor, H. R., Burton, M. J., Haddad, D., West, S., and Wright, H. (2014). Trachoma. The Lancet 384, 2142–2152. doi: 10.1016/S0140-6736(13)62182-0.

Thwaites, T., Nogueira, A. T., Campeotto, I., Silva, A. P., Grieshaber, S. S., and Carabeo, R. A. (2014). The Chlamydia Effector TarP Mimics the Mammalian Leucine-Aspartic Acid Motif of Paxillin to Subvert the Focal Adhesion Kinase during Invasion. J. Biol. Chem. 289, 30426–30442. doi: 10.1074/jbc.M114.604876.

Zagurovskaya, M., Shareef, M. M., Das, A., Reeves, A., Gupta, S., Sudol, M., et al. (2009). EGR-1 forms a complex with YAP-1 and upregulates Bax expression in irradiated prostate carcinoma cells. Oncogene 28, 1121–1131. doi: 10.1038/onc.2008.461.

Zanconato, F., Forcato, M., Battilana, G., Azzolin, L., Quaranta, E., Bodega, B., et al. (2015). Genome-wide association between YAP/TAZ/TEAD and AP-1 at enhancers drives oncogenic growth. Nat Cell Biol 17, 1218–1227. doi: 10.1038/ncb3216.

Zhao, B., Wei, X., Li, W., Udan, R. S., Yang, Q., Kim, J., et al. (2007). Inactivation of YAP oncoprotein by the Hippo pathway is involved in cell contact inhibition and tissue growth control. Genes Dev. 21, 2747–2761. doi: 10.1101/gad.1602907.

