## Supplementary Material for "*Chlamydia trachomatis* induces the transcriptional activity of host YAP in a Hippo-independent fashion"

### 1 Supplementary Data

**Supplementary Data S1.** Summary of differentially expressed genes detected in bulk RNA-sequencing of mock- and Ct serovar L2-infected End1/E6E7 immortalized epithelial cells (End1s). All fold-changes expressed relative to mock-infected control. See enclosed file: “Supplementary Data S1.xlsx”.

**Supplementary Data S2.** Summary of the cross-referencing of Table S1 data with the ChIP Enrichment Analysis (ChEA) database of transcription factor target genes. Individual target list mappings for each transcription factor are available on request. See enclosed file: “Supplementary Data S2.xlsx”.

**Supplementary Data S3.** Summary of differentially expressed genes detected in bulk RNA-sequencing of mock- and Ct serovar L2-infected primary human cervical epithelial cells (HCECs). All fold-changes expressed relative to mock-infected control. See enclosed file: “Supplementary Data S3.xlsx”.

#### 1.1 Supplementary Figures

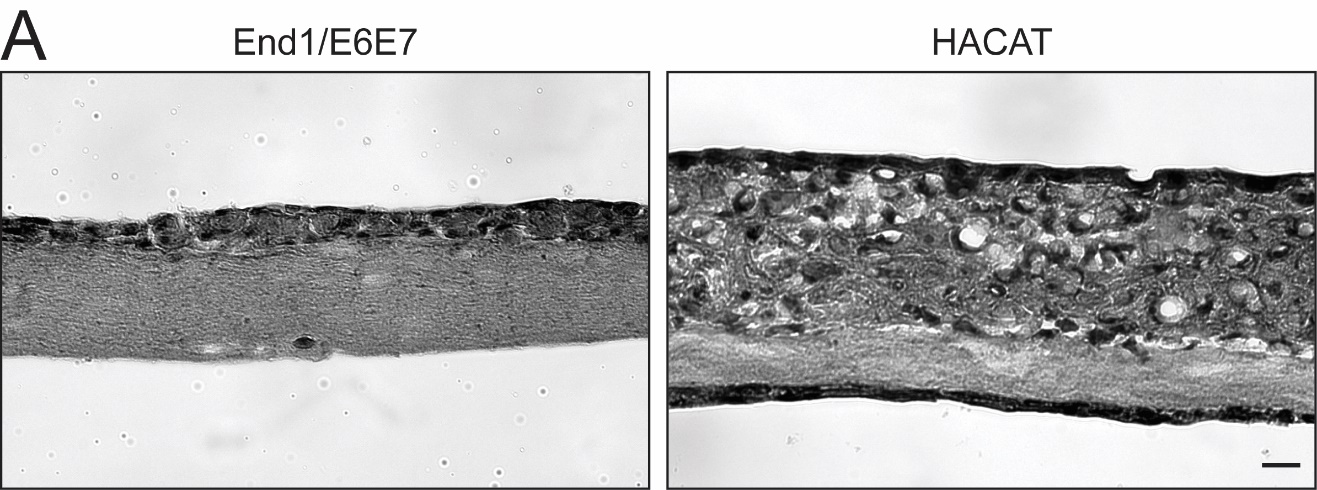

**Supplementary Figure S1.** End1 E6/E7-immortalized epithelial cells in organotypic culture do not exhibit morphology consistent with malignant HPV-associated transformation.
(A) Greyscale, bright-field micrograph of hematoxylin/eosin-stained 20 μm sections of End1 (left) and HaCaT (right) RAFT cultures. Scale bar: 20 μm.

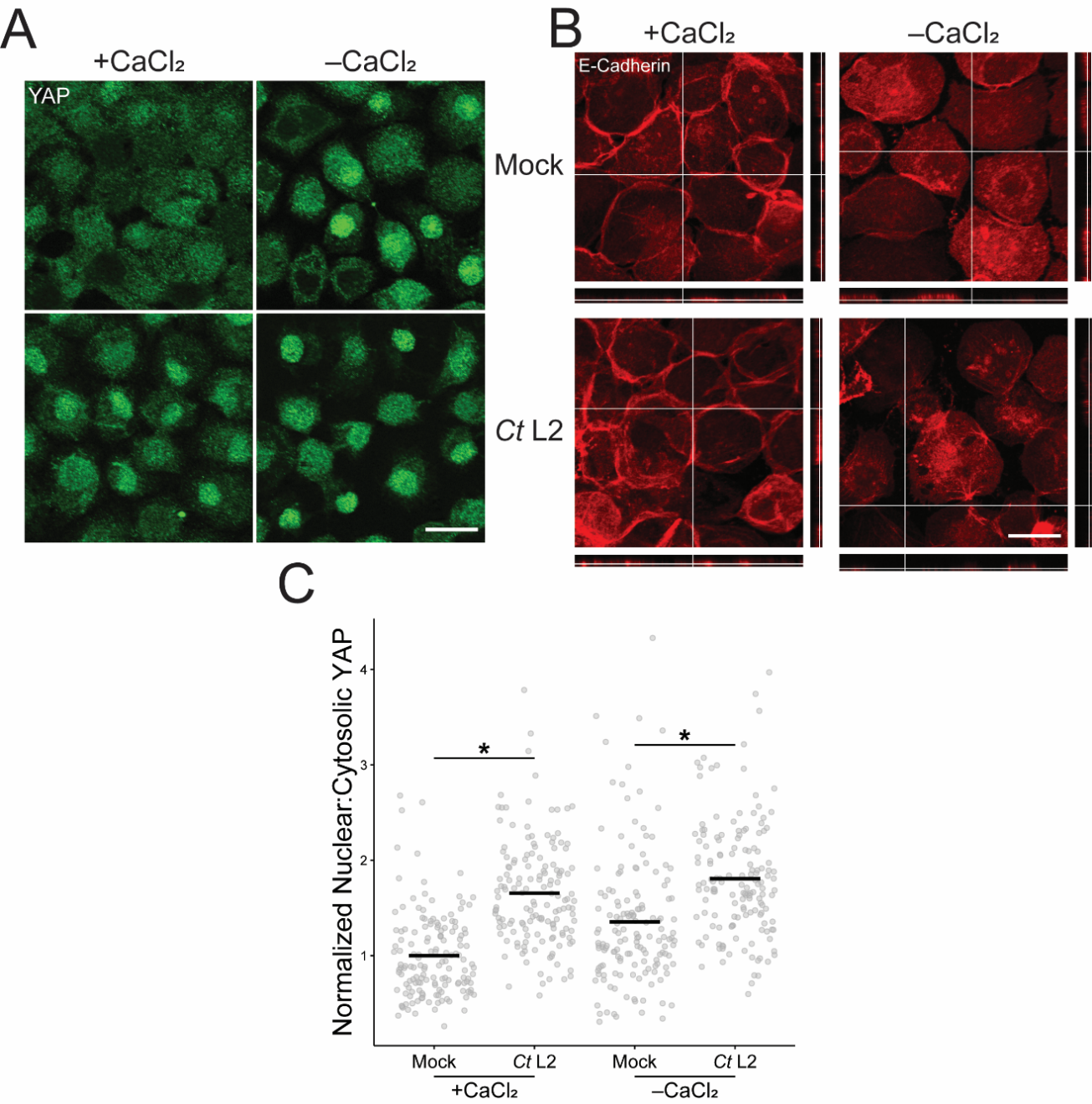

**Supplementary Figure S2.** *Chlamydia* infection enhances YAP nuclear translocation in the Hippo-attenuated condition of adherens junction disruption.
(A) Representative micrographs of YAP (green) translocation at 24 hpi in confluent mock- and *Ct* L2-infected End1 cells cultured in calcium-replete (+CaCl_2_, 0.4 mM) or calcium-deplete (-CaCl_2_) keratinocyte serum-free media. Scale bar: 20 μm.
(B) Representative micrographs of E-cadherin (red) localization at 24 hpi in confluent mock- and *Ct* L2-infected End1 cells cultured in calcium-replete (+CaCl_2_, 0.4 mM) or calcium-deplete (-CaCl_2_) keratinocyte serum-free media. Right sidebar: ZY-plane orthogonal view; left sidebar: XZ-plane orthogonal view; scale bar: 20 μm.
(C) Quantification of YAP nuclear translocation in (A). n = 3 biological replicates, 50 cells measured per sample. Black bars: group means; asterisks: p-values ≤ 0.05, using pairwise Wilcoxon rank sum tests and Bonferroni’s correction for multiple comparisons.
